# Drying conditions alter the defensive function of seed mucilage against granivores

**DOI:** 10.1101/2022.08.15.504038

**Authors:** Eric F. LoPresti, Madison E. Stessman, Sara E. Warren, Katherine Toll

## Abstract

1. Environmental conditions alter the function of many plant traits that drive species interactions, producing context-dependency in the outcomes of those interactions. Seed mucilage is a common, convergently-evolved trait found in thousands of plant species. When wetted, the seed coat swells into a viscid mass; when dried, the mucilage strands strongly cement the seed to whatever it is in contact with.
2. This binding to the ground has been previously shown to protect seeds from granivory. Previous research found both that mucilage volume – and the correlated attachment strength – are higher in species from hot, dry, areas suggesting an environmental component of this trait’s function.
3. Here we (1) quantified the effect of temperature on attachment across many species in a lab setting, (2) tested the potential mechanism behind this correlation by accelerating desiccation speed without changing temperature, and (3) tested whether these relationships introduce context dependency of the defensive function of mucilage in the field, using field trials with harvester ants.
4. We found that (1) increasing temperature during mucilage drying strongly reduced the force needed to dislodge seeds for most species, (2) drying time was likely the driving mechanism behind the loss of attachment strength at higher temperatures, not temperature *per se*, (3) seeds attached to substrate during higher temperatures or under accelerated drying conditions were far more susceptible to granivory.
5. ‘Synthesis’ These results show not only the mechanism behind an abiotic modification of a functional trait of seeds, but that this change majorly alters a key interaction contributing to seed survival. These results add to a small, but growing, literature on the importance of seed mucilage in seed survival and demonstrate strong and largely predictable context-dependency in this trait’s defensive function.

## Introduction

The abiotic environment may alter the function of plant traits important in species interactions, leading to context-dependency in the outcome of these interactions (Maron et al 2014). Under one set of conditions, the effect of the trait on the interaction may be very pronounced and under others, far less so. While it is easy to conclude that “community ecology is a mess, with so much contingency that useful generalisations are hard to find” (Lawton 1999), it is indeed likely that species with similar traits will have parallel responses to changes in the abiotic environment. Having a mechanistic knowledge trait functions can make this contingency predictable and even lead to broad ecological and evolutionary insight about trait function and trait evolution. Seed traits, often quantified in a dry state from herbarium specimens or seed envelopes, are usually considered fixed once the seed has matured and because of the slow rates of cellular processes, far less likely to be affected by the environment than vegetative characters. Here, we investigate how the temperature experienced by a wetted mucilaginous seed alters a defensive function and affects seed survival in the presence of granivores.

Variation in expression of a defensive trait across some abiotic gradient may be genetic (Hahn and Maron 2016), plastic, or some combination of the two. Here we are mostly concerned with the plastic component, how local variation in abiotic conditions changes functions of traits. Wagner and Mitchell-Olds (2018) showed that local conditions across common gardens had large effects on defensive glucosinolate concentrations (up to a doubling) for many *Boechera* genotypes. In a warming experiment in natural plant communities, including 132 plants, Descombes et al (2020) demonstrated this treatment changed putatively defensive traits across most species, but that the outcomes of these changes on interactions with herbivores were idiosyncratic with respect to the treatment. These studies nicely demonstrate environmental effects on defensive traits and interactions, yet making sense of this context-dependency requires some amount of mechanistic knowledge: how exactly does the abiotic environment alter the trait’s role in the interaction across many species?

A powerful method for identifying the underlying mechanism of context-dependence is to experimentally manipulate the environment and quantify how it effects traits and species interactions (e.g., McGuire and Agrawal 2005, Mody et al 2009, Pezzola et al 2017). The classic study done by Lincoln and Mooney (1984) found that herbivory on a monkeyflower was far greater in sun than shade. However, none of the hypothesized defensive traits varied concurrently, instead they concluded that adult butterfly oviposition preferences for sunny areas drove the very strong context-dependency in this system. In contrast, Louda and Rodman (1996) found that reduced glucosinolate concentrations in experimentally sun-exposed mustards, coupled with higher insect abundance in sun, led to markedly higher herbivory in higher light areas. Despite both studies finding higher herbivory in the sun, the former study found the context-dependency was due to arthropod behavior, while concentrations of a defensive chemical contributed in the latter, demonstrating the importance of understanding the mechanisms behind altered interactions.

Though often treated simply as a unit of reproduction, each seed is an individual plant. Most seeds never survive to reproduce, with a major source of mortality in many systems being granivores (e.g. Tevis 1958, Bricker et al 2010). Therefore, defensive traits of the seed may be extremely important in determining survival. Diaspore mucilage (hereafter seed mucilage for simplicity) is a very common, multifunctional trait and occurs convergently in thousands of plant species. The mucilage may occur on the outer layers of seedfruit (i.e. mint nutlets), or accessory fruit (i.e. anthocarps of *Mirabilis*). These seeds are unassuming and entirely nonsticky when dry, yet when wetted, the polysaccharide coating swells into a viscid mass (Figure 1, top row); when dried again, the mucilage strands strongly cement the seed to whatever it is in contact with (Figure 1, middle row). This attachment defends the seed from small granivores (Pan et al 2021), sticks to animals for dispersal (Rønsted et al., 2002), and protects the seed from dislodgement (Pan et al 2022), as well as ameliorating the effects of harsh germination environments (Teixiera et al 2020). The chemical composition, volume, structure, and physical properties of the mucilage layer vary greatly across species (Western 2012, Phan et al 2018, Kreitschitz et al 2021a,b). These differences have functional significance – species with larger mucilage volumes generally had resistance to granivores (Pan et al 2021) and surface flow (Pan et al 2022). Despite previous studies demonstrating mucilage properties across species drive defensive differences (LoPresti et al 2019, Pan et al 2021), mucilage is highly affected by environmental factors, but the ecological consequences of those abiotic factors is uncharacterized.

**Figure 1:**
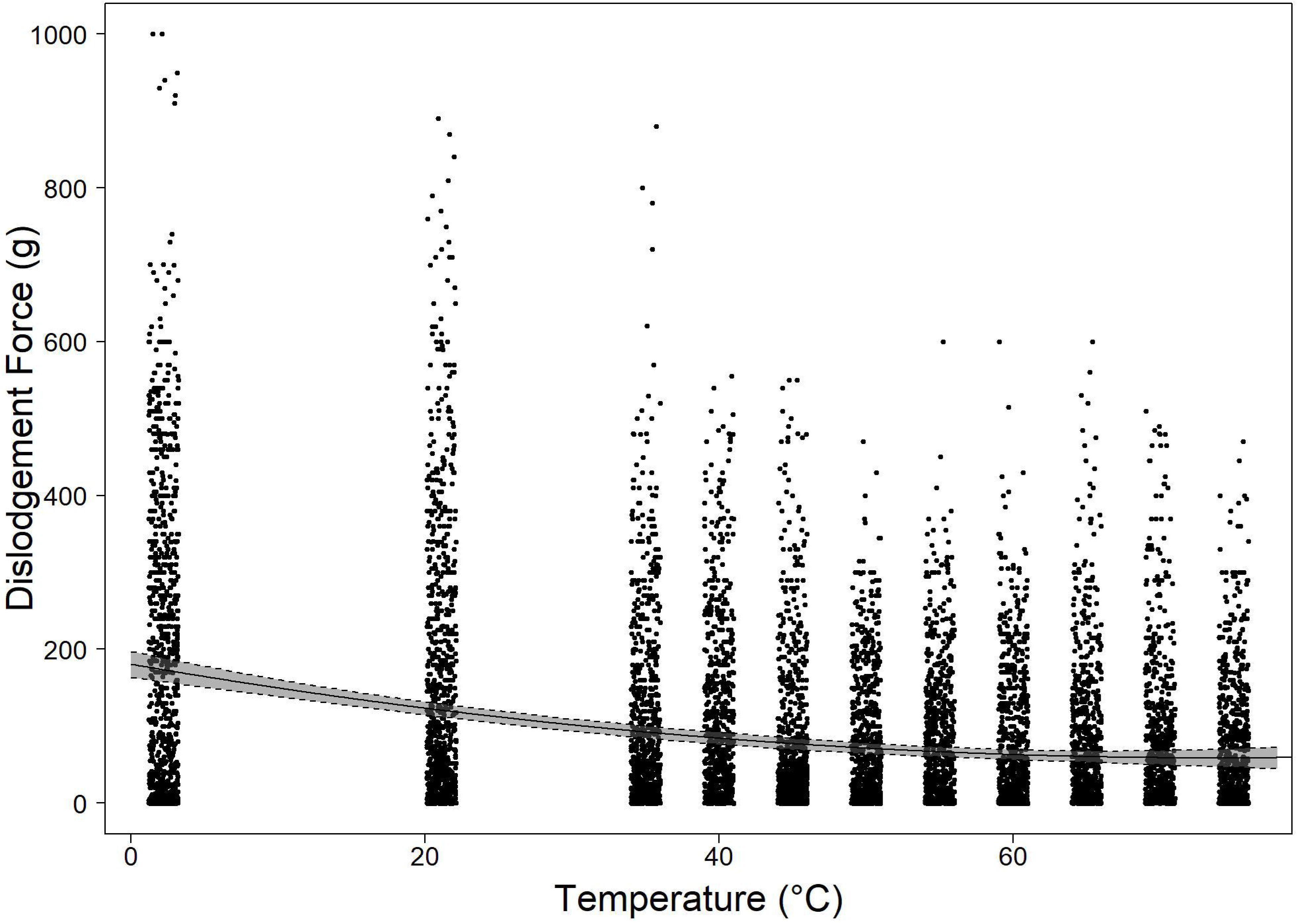
Top row– left to right: mucilage imbibed *Croton captitatus* seed, *Plantago ovata* seed, *Salvia lyrata* nutlet & *Mirabilis nyctaginea* anthocarp. Middle row—left to right: mucilage-bound *Ocimum americanum* nutlet, *Plantago aristata* seed, *Mirabilis nyctaginea* anthocarp, and *Salvia azurea* nutlet. Bottom row—Left: harvester ant clipping mucilage strands of *Plantago rhodosperma*. Right: with visible dried mucilage hanging from a dislodged *Salvia lyrata* nutlet.

Seed mucilage is found in plants in virtually every terrestrial ecosystem worldwide, yet it is thought to be most common and most pronounced in species and populations in hotter, dryer areas and in ruderal plants (e.g. Ellner and Schmida 1981; Al-Shehbaz 1986; Kreitschitz and Valles 2007, Nordenstram et al 2009; Yang et al 2012). While often-stated, only four quantitative comparisons have examined this relationship. Murbeck (1919 in Grubert 1981) found a higher proportion of mucilaginous seeds in the North Africa (xeric) flora than that of Scandinavia (mesic). Ryding (2001) also found that mint species from xeric regions had more mucilage than those in mesic regions. Comparing data from 432 species, Pan et al (2021) found that species from areas of higher temperature, solar radiation, humidity, and fewer wet days per year had higher mucilage attachment potential. Finally, Villellas and Garcia (2012) found that populations of *Plantago coronopus* from areas with lower summer precipitation had higher quantities of mucilage. All four of these interspecific comparisons demonstrated that mucilage evolved more often or to a greater degree in hotter, drier, areas, yet what the effects of climate on mucilage function remain uncharacterized.

Environmental factors clearly affect the functional significance of seed mucilage. In an environment unrealistically entirely devoid of moisture, mucilage will never become imbibed and any functions of imbibition, including defence when mucilage is hydrated and viscous (Yang et al 2013; Pan et al 2021), defence due to mucilage-bound substrate (Fuller and Hay 1983; LoPresti et al 2019), defence due to binding to the ground (Engelbrecht and Garcia-Fayos 2012; Pan et al 2021), or resistance to dislodgement (Engelbrecht et al 2014; Pan et al 2022). The effect of environmental factors on ecological context-dependency in mucilage is entirely unstudied (but see Cowley et al 2022 for a related agricultural study), though Pan et al (2021) cited preliminary data that temperature during drying affects functional properties of the mucilage.

Our study, therefore, seeks to establish how temperature might influence the ecological functions of seed mucilage. We first characterize the effect of temperature on mucilage-binding anchorage across many species. Secondly, we investigate the physical mechanisms behind that relationship. Lastly, we test whether the temperature modification of anchorage strength affects the defensive capabilities of seeds against harvester ants in the field.

## Materials and Methods

### Seed selection

In order to make these results generalizable, we chose 38 species across 10 plant families (Acanthaceae, Brassicaceae, Cistaceae, Euphorbiaceae, Lamiaceae, Linaceae, Nyctaginaceae, Plantaginaceae, Polemoniaceae & Solanaceae). Seeds were obtained commercially or through local collections. The sources of each seed, as well as which species were used in each of the following experiments is detailed in Table S1.

### Drying temperature effects on anchorage strength

To determine if the environmental conditions experienced by the seed during a drying cycle affected anchorage of the seed, we subjected seeds of twenty-two species (Table S1) to a series of temperatures during mucilage drying and then measured the force needed to dislodge them. These protocols are based on those in Pan et al (2021), the measurements from which were found to be ecologically-informative. Seeds of each species were wetted for an hour in order for mucilage to fully imbibe. Ten seeds were then placed onto each of 55 new glass slides (for a total of up to 550 seeds/species tested); washed slides have a lower dislodgement force (Supplementary Figure 3), therefore new slides were used for each trial. The slides were placed in dehydrators (Ivation, 1000W, 10 tray) and set for nine different temperatures, 5 slides per temperature/species. They were dried for an hour, sufficient for full drying. The other two treatments were a room temperature control (5 slides/species), and in the refrigerator (5 slides/species). Even though the dehydrator slides all took <1 hour to dry, all seeds were left at least overnight to dry before measuring dislodgement force. Dislodgement force was measured exactly as in Pan et al (2022; a modification of Pan et al [2021)); briefly, the seeds were dislodged with a shear force applied perpendicularly with a Pesola pressure meter (300, 600, 1000 grams, depending on species) and the force at which dislodgement occurred was recorded. In these trials, we used 10,000 seeds of the 22 species, though one measurement was recorded in error, leaving a final sample of 9,999 dislodgement force measurements.

To analyze these data, we used a mixed-model with a gaussian error distribution, with temperature as a fixed effect and random effect of slide ID nested within species. Given that the response to temperature may be nonlinear, we used a linear term (temperature), a quadratic term (temperature2), and an asymptotic term (1/temperature). We tested models with each combination of these predictors and a no-predictor null model. We also fit a model with each nonlinear predictor for each species independently; the output from these models were plotted to show the individual species relationships (for Supplementary Figure 1), but only the overall model with species as a random effect (as described earlier) is reported.

### Accelerated drying via vacuum desiccator, glass slides

The previous experiment strongly demonstrated that the drying environment temperature strongly influences how tightly the seed attaches during mucilage drying. However, mechanistically, while changing temperature could change the attachment strength directly, it also affects drying speed - at higher temperatures, seeds often appeared fully dry in <10 minutes, whereas the room temperature trials often took 5+ hours for the seeds to appear dry (and they were kept overnight to guarantee that by the time of testing). In order to parse out whether the broad reductions in dislodgement force at higher temperatures were due to this accelerated drying time or due to temperature effects directly, perhaps on mucilage structure, we manipulated these two variables independently. To do this, we dried seeds onto glass slides in two treatments. For each species, we prepared 5 slides, each with 12 seeds, in each of the following ways. Treatment 1 was a control, with seeds left to dry in the lab overnight (as done in the prior experiment, for the 20C treatment). Treatment 2 was a faster desiccation, without raising the temperature (for a comparison with the temperature results). For this treatment, slides of seeds were placed into a vacuum desiccator that contained drierrite, a desiccant. For these trials, we tested 1181 seeds of 10 species.

### Accelerated drying on sandpaper, via heating and vacuum desiccator

Additionally, we manipulated the substrate, and ran an additional test with those two treatments, plus a third (a 75C heat treatment, as in the prior section) on glass slides covered in 80-grit sandpaper (3M - 80 grit), as used in the ant trials. As in the other treatments, these were dried overnight, though actual drying time for the latter two treatments, visually, was less than an hour. During drying, humidity in the vacuum chambers was 18%-21%, oven 10-18%, and lab 45%-65%. The force needed to dislodge was measured for these seeds in the same manner as previously. All species used in ant trials were planned to be tested in this way in order to quantitatively link this reduction to field performance, but the vacuum went out in the building before all could be tested. In these trials, we test 1725 seed of 10 species.

For both the glass slide test and the sandpaper tests, we analyzed the overall treatment effect on dislodgement force, using a mixed-model with a guassian error distribution and treatment as a fixed effect, with species as a random effect (adding date as a random effect did not improve the fit). In order to calculate post-hoc comparisons within species, we did a linear regression of dislodgement force as a function of the interaction between treatment and species, calculating posthoc comparisons of treatment within species using a Tukey test in the R package *emmeans*.

### Seed defence trials

To test whether the relationships established in the first two parts of this study translated into a context-dependent interaction with granivores, we did granivory trials using the common harvester ant, *Pogonomyrmex barbatus*. These experiments occurred along gravel paths and mown lawns at the Oklahoma State University Botanical Garden, in Stillwater, OK, where *P. barbatus* is common. Seven nearby nests were selected for use in this experiment. Seed treatments were prepared as in the previous experiment (control, vacuum desiccated, heat dried) with one important difference. Instead of being attached directly to glass slides, the glass slides again had a layer of sandpaper (3M - 80 grit) glued on, which is a more appropriate substrate proxy than glass for ant foraging. A trial consisted of three slides (containing 5-12 seeds, depending on seed size), one of each treatment, that were placed in a semicircle ~0.5m from the entrance of the nest. As in LoPresti et al (2019), these were monitored until one treatment was entirely consumed, or for two or four hours, depending on the ant activity (i.e. some days, probably driven by colder weather, very few seeds would be consumed in two hours and those trials were lengthened to four hours). Multiple trials of different species were done at the same nests simultaneously, but a species was never repeated at the same nest on the same day. Two species (*Nicandra physalodes* & *Salvia sclarea*) had no vacuum treatment, only the heat and control treatments. The sample size was 7699 seeds in total from 34 species.

We analyzed the ant trial results using seed survival as a binomial response variable (survived or was removed) with treatment as a fixed effect and random effects of date, ant nest, species, as well as a composite ID for each slide nested within one for each species trial (i.e., *Camelina sativa* at ant nest #3 on July 7^th^). We compared treatment means using a Tukey’s posthoc comparison of means.

### Data and analyses

All data and code is available on FigShare (doi: data: 10.6084/m9.figshare.20477151.v2, temperature analysis script: 10.6084/m9.figshare.20477142.v2 rest of analyses script: 10.6084/m9.figshare.20477145.v2, graphing script: 10.6084/m9.figshare.21543009.v1). All analyses were conducted and figures made in R vers. 4.1.0. Specific model parameters are described in the sections above.

## Results

### Higher drying temperatures reduce anchorage strength

Generally, higher drying temperatures greatly reduced the amount of force necessary to dislodge the seeds (Figure 2; Supplementary Figure 1). A linear mixed-model with drying temperature as linear, exponential, and asymptotic fixed effects and species as a random intercept (i.e. Force ~ Temp + Temp2 + 1/Temp + [1|Species]) fit better than any simpler model and each of these coefficients was highly significant (Likelihood ratio tests, p< 0.01 for each). However, individual species responses varied greatly (Supplementary Figure 1). The effect of temperature was especially pronounced in the plantains (*Plantago* spp.) and flaxes (*Linum* spp.). The highest temperature treatment for *P. aristata* required only 1.5% the force of the lowest temperature treatment for dislodgement! *P. rhodosperma* (5.5%), and *P. virginica* (8.7%) were similar, with *P. patagonica* having a slightly smaller effect (26.2%). The highest temperature for *Linum lewisii* had a mean dislodgement force of just 2.8% of that at the lowest temperature, with *L. grandiflorum* at 13.3%, *L. usitassimum* at 19.1%, and *L. perenne* at 30.0%. In contrast, other groups were far less affected; certain species showed no significant decrease in dislodgement force at higher temperatures (i.e. *Helianthemum nummularium*). Nevertheless, on average seeds dried at the highest temperatures required ~33% of the force to dislodge compared to those dried in the refrigerator or at room temperature, demonstrating that attachment is strongly dependent on temperature.

**Figure 2:**
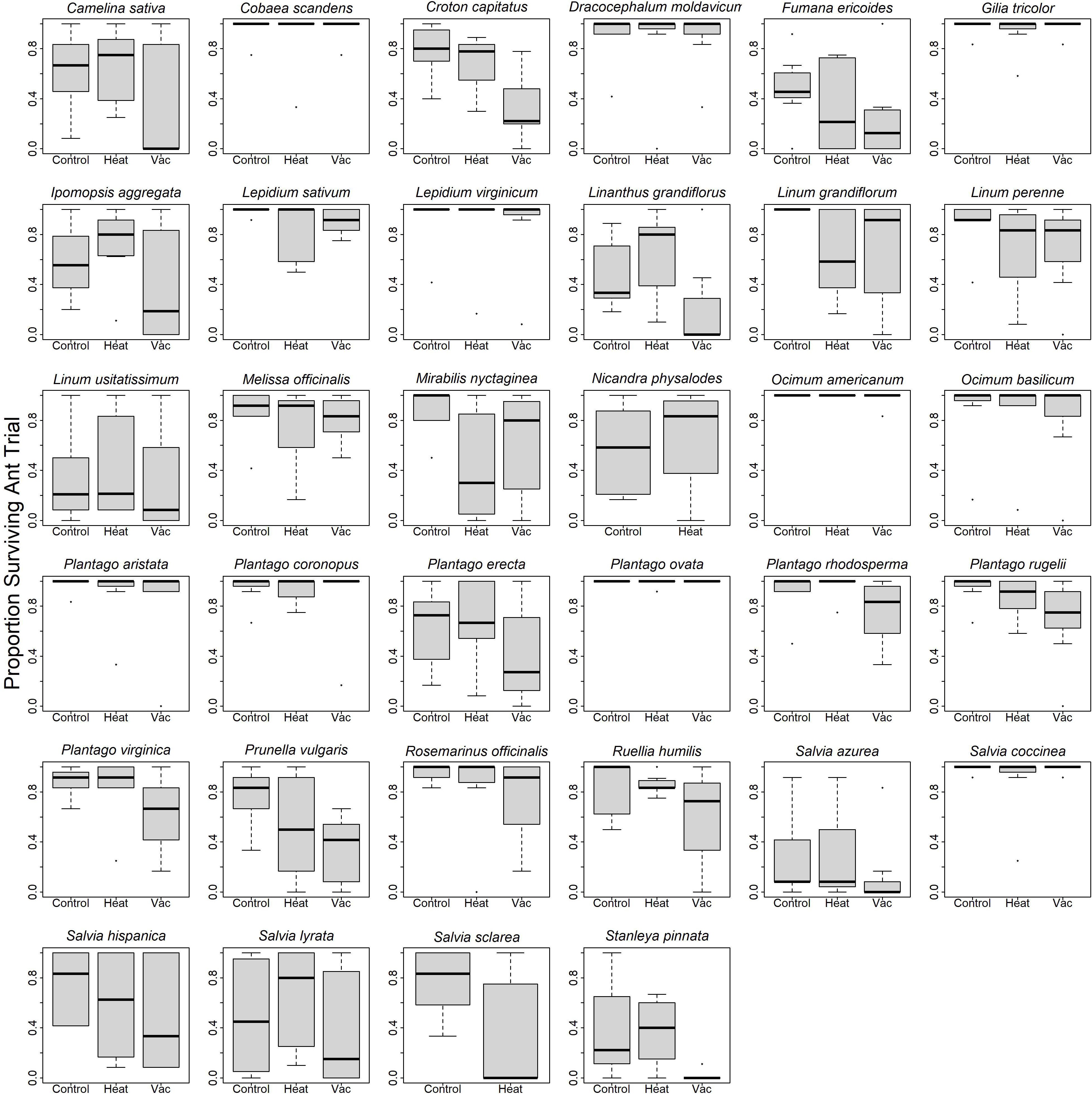
Overall relationship between drying temperature and force necessary to dislodge a seed from a new glass slide for the 16 species tested. Predictions from the best-fitting linear mixed-model are plotted; species was treated as a random effect, so these results may be thought of as the “average” species. For plots of each individual species’ response, see Supplementary Figure 1.

### Humidity manipulations to drying speed, independent of temperature

#### Accelerated drying via vacuum desiccator on glass slides

Using a vacuum desiccator greatly accelerated drying speed while remaining at room temperature, isolating the effect of drying speed from temperature (Figure 3). Seeds in the vacuum desiccated treatment required significantly less force to dislodge than those at room temperature overall (Mixed-effect linear model, with species as a random effect, post-hoc comparison of treatment means, p < 0.001; Figure 3), mirroring the results of higher temperatures. Post-hoc comparisons of treatment means were significant for six of the ten species tested and the means of the control groups were higher than the vacuum for nine of the ten species tested (Figure 3).

**Figure 3:**
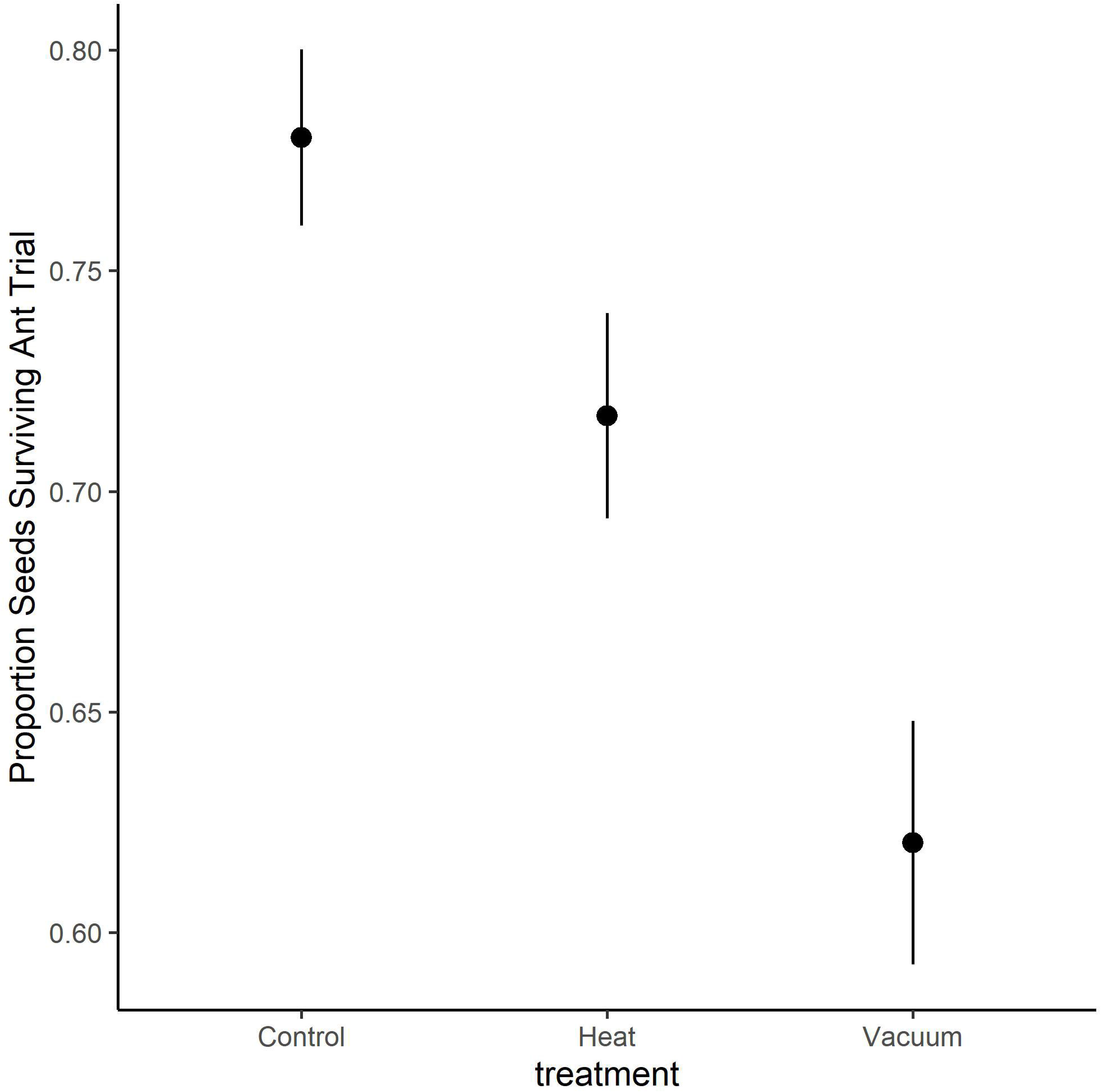
Dislodgement force necessary to dislodge seeds dried to glass slides at room temperature (room) and in vacuum desiccators with Drierite (vac). Shading indicates significant (p<0.05) post-hoc comparison of means for each treatment/species combination, from the best-fitting interactive (treatment*species) model.

#### Accelerated drying via vacuum desiccator and heating on sandpaper

Accelerated drying, via heat and vacuum dehydration, greatly reduced the force needed to dislodge the seed (Mixed-effect linear model, with species as a random effect, posthoc comparison of treatment means, p < 0.001; Figure 4). The control group had a higher mean dislodgement force than the heat and vacuum treatments in all of the species tested; post-hoc comparisons the control group showed it differed significantly from the heat treatment in seven of ten species, the same number (and same species) differing in control versus vacuum treatments.

**Figure 4:**
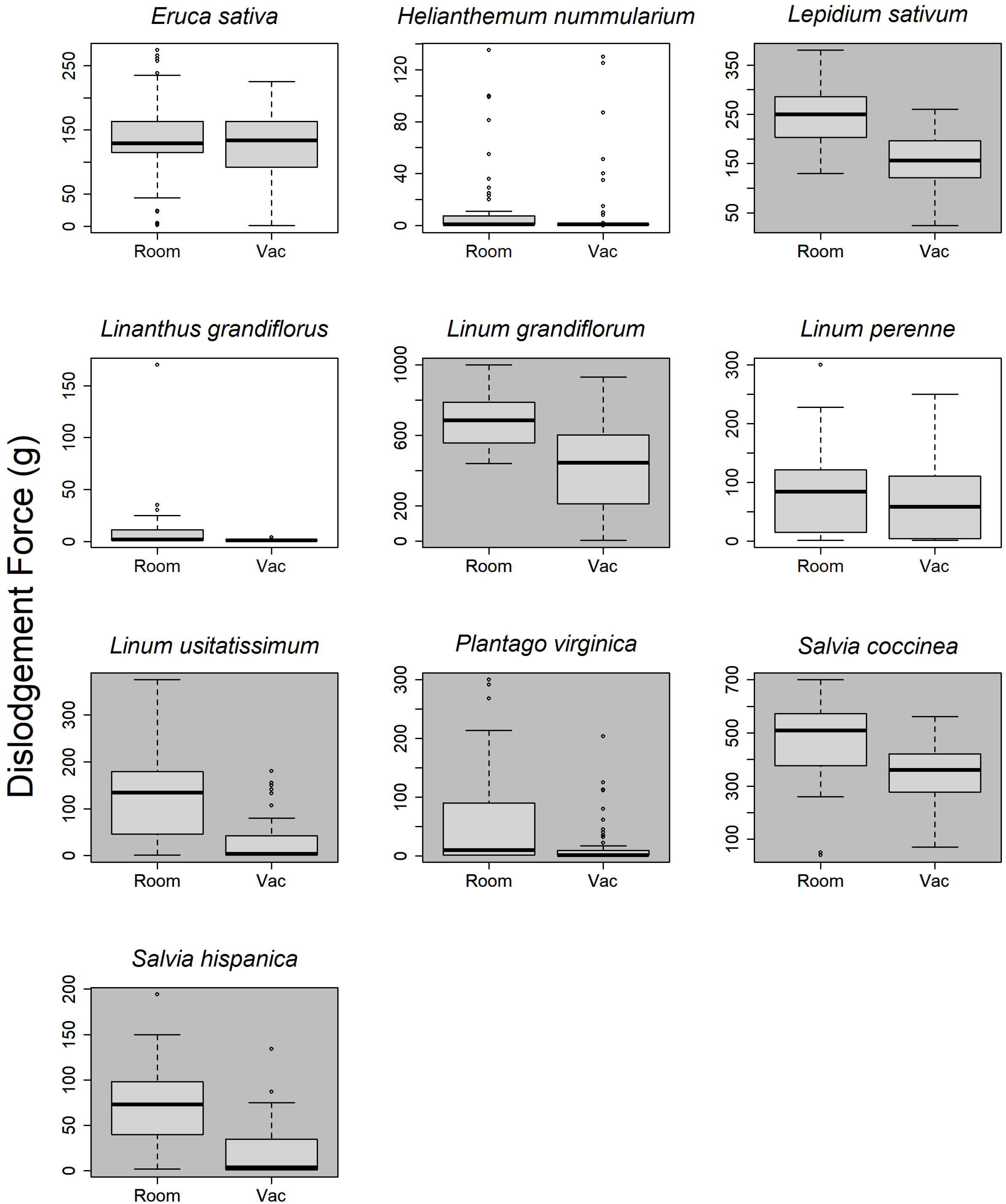
Mean force necessary to dislodge seeds dried to sandpaper at room temperature (Control), in a drying oven at 70C (HighTemp), and in vacuum desiccators with Drierite (Vac). Similar shading within a plot indicates no difference in a post-hoc comparison from a linear model with treatment*species as fixed effects (two species where shading is not obvious. Both were not significant and denoted as “n.s.”).

### Functional role of attachment drying speed in defence against granivory

Accelerated drying due to heat or vacuum increased granivory on seeds (Figure 5) compared to the control group (Tukey’s post-hoc comparison of means, all pairwise comparisons p < 0.01). Seeds dried in the vacuum desiccator were also removed at a significantly higher rate than those in the heat-dried treatment. This effect, however, was not uniform – some were removed nearly completely, others barely removed, and the treatment effects differed depending on the species. The overall effect, with species as one of the random effects, was strongly significant, yet it may be useful to examine the species separately to determine the generality of this result (supplementary figure 2). While only 56% of the species (19/34) showed a significant reduction in survival in the heat or vacuum compared to the control (post-hoc comparisons in an interactive model), 68% (23/34) had a higher mean survival in control than heat, with only 29% lower survival (10/34, 1 had no difference). This was a significant result using a sign-test (p=0.035). Mirroring the overall results, the difference was even stronger between the control and vacuum; 91% (29/32) of species had higher survival of controls than vacuum-dried, with the remaining 9% exactly equal (3/32), a highly significant difference with a sign test (p<0.001).

**Figure 5:**
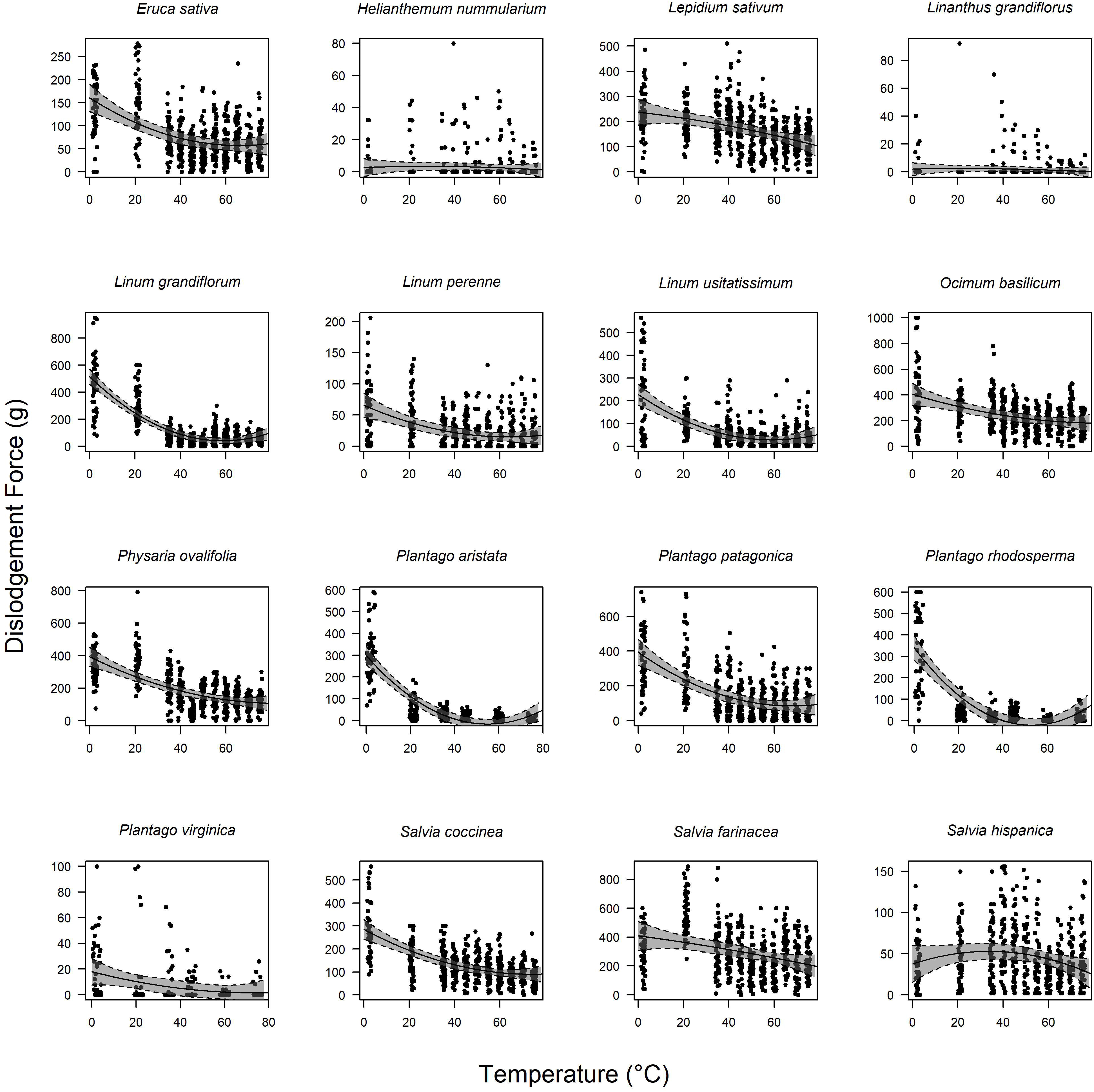
Survival of seeds exposed to harvester ants, when dried to dried to sandpaper at room temperature (Control), in a drying oven at 70C (Heat) and in vacuum desiccators with Drierite (Vacuum). Dot is the predicted mean, lines indicate 95% CI of the mean. Each treatment differed significantly from each other in a post-hoc comparison of means. For plots of individual species responses, see supplementary figure 2.

## Discussion

Environmental conditions modified the ecological function of an important seed trait – higher drying temperatures led to lower seed survival in the presence of harvester ants, due to easier removal of seeds. As hypothesized, this context dependency was largely predictable across species, though the mucilage traits differ greatly across these many evolutionary origins (Grubert 1974, Kreitschitz et al 2021b; Pan et al 2021). While not every species showed a significant reduction in dislodgement force at higher temperatures or faster drying speeds, most species had a measurable reduction. Similarly, the reduction in defensive efficiency was rather uniform for those species palatable to the ants, with few species having either treatment with lower attachment strength (i.e. high temp or vacuum dried) surviving at higher levels than the room control treatment. Unlike a previous study that found an effect of attachment strength in interspecific comparisons (Pan et al 2021), in which other traits surely played a large role, the intraspecific comparisons here are much stronger support for a protective benefit of strength of attachment as we manipulated it within each species. The interplay between this widespread trait - found in 1000’s of plant species (Grubert 1983) - the environment, and interacting organisms is an exciting topic that we hope these results inspire future studies of.

### The environment affects mucilage attachment strength

High temperatures greatly reduced the force required to dislodge seeds. We experimentally demonstrated the mechanisms underlying the modification of the interaction - higher temperatures led to weaker attachment force, via a decrease in drying time. While these two factors are mechanistic at some level, this is not a full physical explanation for how this reduction occurs - that would require further study. However, we have a strong mechanistic hypothesis integrating these results and previous research. In interspecific comparisons of over fifty species, attachment strength was strongly correlated with mucilage volume (Pan et al 2021, 2022). Mucilage volume probably affects attachment strength by altering the area of contact (hypothesized in Grubert 1974); therefore, it is likely that the rapid decline in volume at higher temperatures or at low humidity simply led to a smaller area of substrate that the mucilage strands were in contact with; it is likely that the reduced attachment strength in several species in the refrigerator treatment was due to the humidity control in household refrigerators. For this reason, we can also hypothesize that other environmental factors that affect mucilage volume (such as pH, ion concentrations, number of wetting-drying cycles: Grubert 1974, Muñoz et al 2012) or area that the mucilage contacts (i.e., seed size or substrate grit, Stessman unpublished data) would similarly alter interactions and subsequent seed success. Examining individual seed attachment area and dislodgement force, would be a straightforward, if labor-intensive way, of confirming this mechanistic hypothesis.

While most plants cannot survive very high temperatures, substrate surfaces, where many seeds are, reach the highest treatments we applied (Nobel et al 1986, pers. obs.). While it is unlikely that a seed would be imbibed at that temperature (as any substrate reaching that temperature requires abundant sunshine), it is likely that many of the higher temperatures we used are regularly encountered by seeds and harvester ants (Whitford and Ettershank 1975) and that this fundamental interaction is therefore likely to be greatly modified by temperature in realistic systems. Of course, temperature itself was not the ultimate driver, and while seeds are rather unlikely to encounter vacuum chambers of desiccant in the real world, realistic differences in humidity are also certainly likely to modify these interactions as they would alter drying speed.

Temperature may affect other aspects of the functional ecology of seed mucilage besides granivory. Few studies have examined attachment strength of mucilage (but see Grubert 1974, Kreitschitz et al 2021, Pan et al 2021) and none have examined the effect of temperature, but it is well-known that other seed mucilage properties change with temperature in its food science, pharmaceutical and industrial applications. Temperature-dependent changes in physical properties of mucilage have been characterized for economically-important species used in this study (i.e. flax, *Linum usitatisimum*: Mazza and Biliaderis 1989, chia, *Salvia hispanica*: Muñoz et al 2012, and arugula, *Eruca sativa*, Koocheki et al 2012). However, few studies have examined the effects of temperature on ecologically-important processes involving seed mucilage. The two studies that have both concern germination, a process that mucilage may either promote or retard (Western 2012). Gorai et al (2014) examined germination of *Henophyton deserti* under seven temperature treatments and found germination differences between mucilage intact and demucilaged seeds under only one temperature. Similarly, Bhatt et al (2016) found a significant effect of the interaction between temperature and mucilage presence in two of five species tested (*Lavandula subnuda* and *Lepidium aucheri*). Given the diverse roles of mucilage (Grubert 1974; Western 2012), it is likely that much further complexity exists and when thinking about mucilage’s roles in the field, the effect of simultaneous functions, including defence and germination, ought to be examined concurrently.

### Reduced strength of attachment affects seed survival

Unsurprisingly, the effect of this reduced attachment strength due to higher temperatures or more rapid drying made the seeds far easier to exploit by ants. This result makes good sense; a seed bound to the ground requires a dislodgement force between 10,000-30,000x its own weight to dislodge (Grubert 1974), and interspecific differences in attachment strength explain protection from ants well (Pan et al 2021). An ant cannot, therefore, simply pick up even the most weakly-bound seeds (i.e. <10 g required to dislodge). Instead, they clip the dried mucilage strands (Figure 1), increasing the time needed to collect and probably greatly increasing the cost of collecting the seed (see supplementary information in Pan et al 2021 for a video of this behavior). It is worth stressing that we were not explicitly testing the defensive benefit of mucilage *per se*, which would have had a non-bound treatment. Previous experiments have found very strong effects of binding on survival across many species (e.g., Gutterman and Shem-Tov 1997, Engelbrecht and Garcia-Fayos 2012, Pan et al 2021), and ants certainly struggled and gave up on tightly bound seeds in this experiment (LoPresti, Stessman, pers. obs.) and therefore the defensive function was assumed, and here we investigated only the context-dependency of the benefit.

Attachment strength, however, is far from the only defensive trait that seeds have. The drying environment had little effect on survival of a few species in the study (Supplementary Figure 2), most notably those species that were not collected at all (*Ocimum americanum* & *Plantago ovata*), or collected in extremely small numbers (*Cobaea scandens*, *Lepidium densiflorum, Gilia tricolor, Plantago aristata*, & *Salvia coccinia*). For *C. scandens*, the seeds were simply too large to be easily exploited, but for others, the ants simply rejected them after antennating or palpating them with their mandibles (pers. obs.). It is possible - even likely - that the mucilage layer plays a role in this advertisement of unpalatability as well. It is both the outer layer of the seed, which is first contacted by granivores and therefore, one that may deter them (see LoPresti 2017). In fact, the diversity of secondary chemistry in mucilage (e.g., Wardle et al 1991), could be an advertisement of unpalatability. Mucilage also protects seeds when imbibed (Yang et al 2013, LoPresti et al 2019), and when it binds substrate particles to the seed (Fuller and Hay 1983, LoPresti et al 2019). Therefore, mucilage may play a greater role in defence than simply attachment strength and the chemistry and other physical aspects of this layer deserve greater attention as a multifunctional defensive trait.

### What is the environment like when seeds become imbibed?

Many mucilaginous seeds simply drop off plants upon dehiscion (e.g. Fuller and Hay 1983), though there may be some plant control of when this occurs. *Cardamine hirsuta* seeds are ballistically dispersed, and while Vaughn et al (2011) suggest that herbivore touch stimulates them in natural populations, our observations following rain events in Oklahoma, US, suggest that many siliques rupture and the seeds disperse during rain storms. These highly mucilaginous seeds are thus imbibed in the wet environment and often cemented to surrounding vegetation or objects (tree trunks, fallen wood, the ground, e.g. Figure 6). Rain may be a common mechanism for mucilaginous seeds to disperse from a plant, other examples include the dispersal of many Aizoaceae, *Thlaspi*, many Lamiaceae, *Collomia*, and others (Parolin 2005, LoPresti, pers. obs.). As these seeds leave the plants during wet periods, the mucilage begins imbibing quickly (e.g. Yang et al 2013) and the seeds get firmly stuck to a substrate during this initial dispersal period.

**Figure 6:**
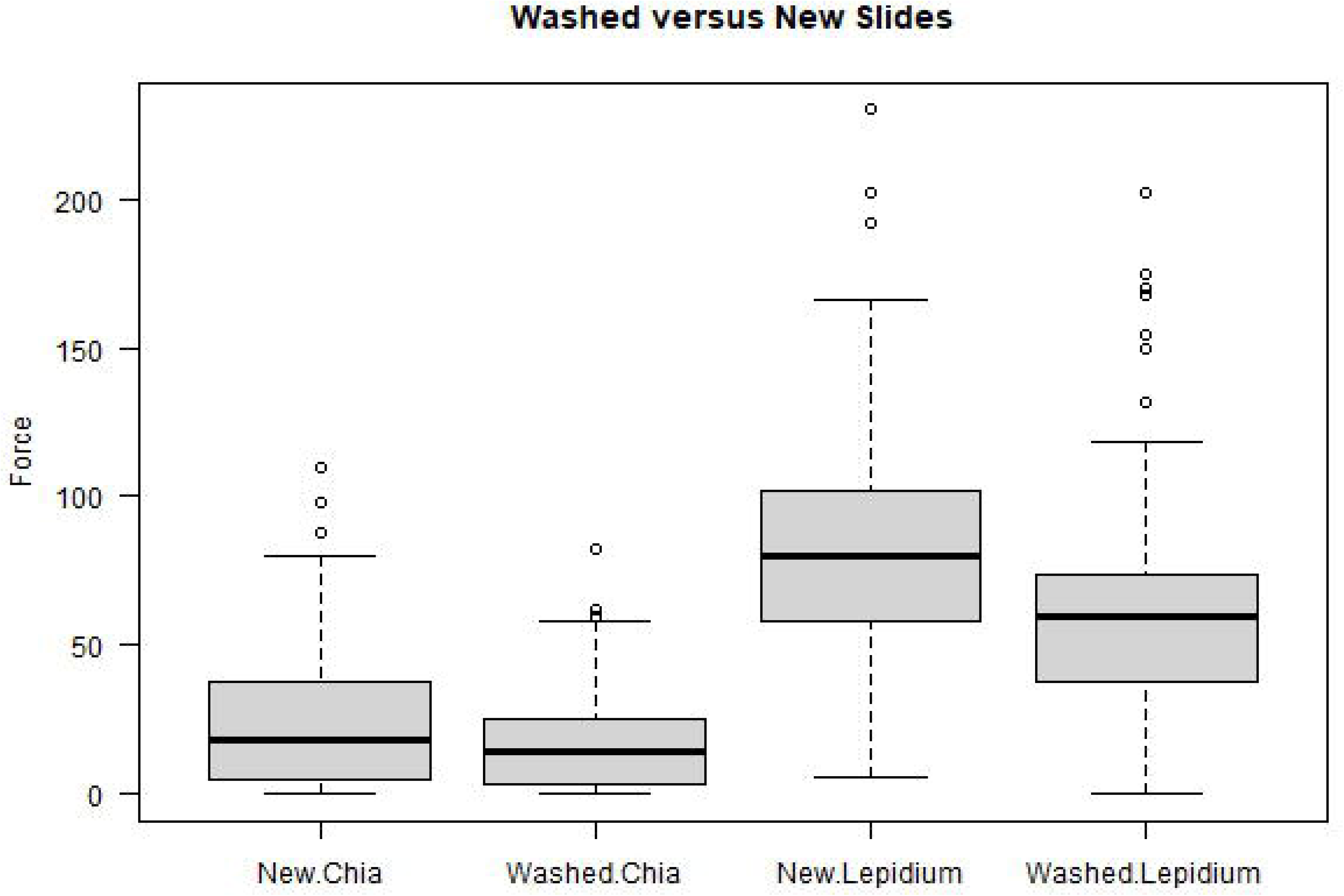
*Plantago rhodosperma* seeds naturally mucilage-bound to a rock, 6-July-2022. The most recent precipitation was ~1 cm, 10-June, therefore, these must have remaining attached for at least that long, if not longer. Boomer Lake Park, Stillwater, Oklahoma, USA. Photo: KT

Despite repeated wetting-drying periods due to precipitation and even morning dew (Grubert 1974, Gutterman and Shem-Tov 1996, Huang et al 2008), those seeds that attach at that point are unlikely to move far. While mucilage is rehydratable, seeds attached to a substrate are not easily dislodged. The exact mechanism for this phenomenon deserves further study; Pan et al (2022) found that even after a week of being under constant water flow, most previously dried seeds remained firmly stuck to tiles. This unexpected result, coupled with observations of mucilaginous seeds firmly attached to substrates and objects naturally, demonstrates that the attachment of these seeds is both ecologically realistic and that the conditions that they experience during that initial drying period probably determine their subsequent success.

### Do plants that regularly experience high temperatures evolve to have more mucilage and stronger attachment?

Previous research found that seed mucilage of species from warmer areas had stronger attachment and greater mucilage volume than that of species from cooler areas (Pan et al 2021, also see Villelas and Garcia 2012 for population level differences). Coupled with the findings in this study of stronger attachment at lower drying temperatures, we think it is reasonable to assume that species or populations regularly experiencing higher temperatures may need greater attachment strength to obtain the same protection that may come with the rapid drying in warmer areas. Therefore, we believe that in addition to the environment modifying the function of mucilage, it also likely is a potent selective force that may, in part, drive the diversity of mucilage properties we see across plants (Grubert 1974, Western 2012, LoPresti et al 2019, Cowley et al 2021; Pan et al 2021, 2022).

While differences in attachment strength are likely to drive differential survival between seeds, it is likely that mucilage volume, as the underlying trait strongly correlating with force needed to dislodge the seed (Pan et al 2021, 2022) is the heritable component that selection would act on. Mucilage volume differs greatly among species, populations and between individuals, likely indicating some degree of heritability (e.g. Villelas and Garcia 2012; Teixiera et al 2020; Cowley et al 2021; Pan et al 2021,2022). Population examinations, especially of widespread species spanning a large climatic range, (e.g. Villelas and Garcia 2012), or examinations of a complete clade would be the strongest tests of this hypothesis. Fortunately, many mucilaginous-seeded species are nearly cosmopolitan as introductions (e.g., *Capsella bursa-pastoris, Euphorbia prostrata*, *Glechoma heracea, Arabidopsis thaliana, Plantago coronopus, lanceolata, & major, Prunella vulgaris*) or extremely widespread naturally (e.g., *Plantago maritima*) and present rich opportunities to study.

### Future directions

The results of these experiments demonstrate strong context-dependency in the defensive function of seed mucilage, driven by altered drying speed. This totally new line of research brings up more questions than it answers. We suggest that future research investigate:

1. How these temperature-mediated physical changes are affected by different chemical compositions of mucilage and mucilage location (e.g., Phan and Burton 2018, Phan et al 2020).
2. The interplay between attachment strength and area of attachment, with consideration of both mucilage volume and seed shape (i.e., the flat seeds of flax versus the nearly spherical seeds of chia).
3. How much attachment strength affects foraging decisions of ants (i.e., by changing the costs of foraging).
4. Functional consequences of other factors that alter attachment strength (i.e., pH, substrate, wetting-drying cycles) for granivory and other functions of mucilage.
5. How attachment strength and seed survival vary across a season for seeds in real populations, due to dial and seasonal differences in temperature, precipitation, and humidity.

## Acknowledgements

Vincent Pan discovered the temperature and dislodgment force relationship and encouraged us to pursue this study. We thank Gabby Barber, Addison Darby, Michael Foisy, Sierra Jaeger, and Lizz Waring for feedback on the study and manuscript. Michelle Dextraze collected the *Plantago rugelii* seeds. SEW was funded by the OSU College of Arts and Sciences AURCA program, with special thanks to Rachel Eaton; supplies and MES were funded by start-up funds to EFL from Oklahoma State University.

## Author Contributions

EFL conceived of the study. SEW conducted the temperature dependence trials. MS conducted the vacuum and sandpaper trials. MES, EFL, KT conducted the ant trials. EFL analyzed the data and wrote the manuscript. All authors edited and approved the manuscript.

## Data Availability Statement

Data and scripts are available at Figshare at DOI: 10.6084/m9.figshare.20477151 (full data), 10.6084/m9.figshare.20477145 (granivory and vacuum script), 10.6084/m9.figshare.20477142.v2 (adherence/temperature script), and 10.6084/m9.figshare.21543009.v1 (figure scripts).

## References

Al-Shehbaz, I.A., 1986. New wool-alien Cruciferae (Brassicaceae) in eastern North America: Lepidium and Sisymbrium. Rhodora. 88: 347–355.

Bhatt A., Santo A., Gallacher D. 2016. Seed mucilage effect on water uptake and germination in five species from the hyper-arid Arabian desert. Journal of Arid Environments 128: 73–79.

Bricker M., Pearson, D. & J. Maron. 2010. Small-mammal seed predation limits the recruitment and abundanceof two perennial grassland forbs. Ecology 91:85–92

Cowley, J.M., McNeil, D.L., Lui, K.Y., Barsby, J.P., Ciani, S., Cerne, V. & Burton, R.A., 2022. Rain events at maturity severely impact the seed quality of psyllium (Plantago ovata Forssk.). Journal of Agronomy and Crop Science. 208: 567–581.

Descombes, P., Kergunteuil, A., Glauser, G., Rasmann, S., Pellissier, L., 2020. Plant physical and chemical traits associated with herbivory in situ and under a warming treatment. Journal of Ecology, 108: 733–749.

Ellner S, Shmida A. 1981. Why are adaptations for long-range seed dispersal rare in desert plants? Oecologia 51: 133–144.

Engelbrecht M, Bochet E, García-Fayos P. 2014. Mucilage secretion: an adaptive mechanism to reduce seed removal by soil erosion?: mucilage secretion and seed removal. Biological Journal of the Linnean Society 111: 241–251.

Engelbrecht M, García-Fayos P. 2012. Mucilage secretion by seeds doubles the chance to escape removal by ants. Plant Ecology 213: 1167–1175.

Fuller PJ, Hay ME. 1983. Is glue production by seeds of Salvia columbariae a deterrent to desert granivores? Ecology 64: 960–963.

García-Fayos P, Engelbrecht M, Bochet E. 2013. Post-dispersal seed anchorage to soil in semiarid plant communities, a test of the hypothesis of Ellner and Shmida. Plant Ecology 214: 941–952.

Gorai M, El Aloui W, Yang X, Neffati M. 2014. Toward understanding the ecological role of mucilage in seed germination of a desert shrub Henophyton deserti: Interactive effects of temperature, salinity and osmotic stress. Plant and Soil 374: 727–738.

Grubert M. 1974. Studies on the distribution of myxospermy among seeds and fruits of Angiospermae and its ecological importance. Acta Biologica Venezuelica 8: 315–551.

Grubert M. 1981. Mucilage or gum in seeds and fruits of angiosperms: a review. Munich, Germany: Minerva-Publikation.

Gutterman Y, Shem-Tov S. 1997. The efficiency of the strategy of mucilaginous seeds of some common annuals of the negev adhering to the soil crust to delay collection by ants. Israel Journal of Plant Sciences 45: 317–327.

Hahn, P.G., Maron, J.L., 2016. A framework for predicting intraspecific variation in plant defense. Trends in Ecology & Evolution. 31: 646–656.

Huang Z, Boubriak I, Osborne DJ, Dong M, Gutterman Y. 2008. Possible role of pectin-containing mucilage and dew in repairing embryo DNA of seeds adapted to desert conditions. Annals of Botany 101: 277–283. doi:10.1093/aob/mcm089.

Koocheki, A., Razavi, S. and Hesarinejad, M.A., 2012. Effect of extraction procedures on functional properties of Eruca sativa seed mucilage. Food Biophysics 7: 84–92.

Kreitschitz A, Kovalev A, Gorb SN. 2015. Slipping vs sticking: Water-dependent adhesive and frictional properties of Linum usitatissimum L. seed mucilaginous envelope and its biological significance. Acta Biomaterialia 17: 152–159.

Kreitschitz A, Haase E, Gorb SN. 2021a. The role of mucilage envelope in the endozoochory of selected plant taxa. The Science of Nature 108: 2.

Kreitschitz A, Kovalev A, Gorb SN. 2021b. Plant seed mucilage as a glue: Adhesive properties of hydrated and dried-in-contact seed mucilage of five plant species. International Journal of Molecular Sciences 22: 1443.

Kreitschitz, A. and Vallès, J., 2007. Achene morphology and slime structure in some taxa of Artemisia L. and Neopallasia L.(Asteraceae). Flora-Morphology, Distribution, Functional Ecology of Plants 202: 570–580.

Lawton J.H. 1999. Are there general laws in ecology? Oikos 84: 177–192

Lincoln, D.E. and Mooney, H.A., 1984. Herbivory on Diplacus aurantiacus shrubs in sun and shade. Oecologia, 64: 173–176.

LoPresti EF. 2016. Chemicals on plant surfaces as a heretofore unrecognized, but ecologically informative, class for investigations into plant defence. Biological Reviews 91: 1102–1117.

LoPresti EF, Pan V, Goidell J, Weber MG, Karban R. 2019. Mucilage-bound sand reduces seed predation by ants but not by reducing apparency: a field test of 53 plant species. Ecology 100: e02809.

Louda, S. M, Rodman, J.E. (1996). Insect herbivory as a major factor in shade distribution of a native crucifer (*Cardamine cordifolia* A. Gray, Bittercress). Journal of Ecology, 84: 229–237.

Mazza, G, Biliaderis, C.G. (1989). Functional properties of flax seed mucilage. Journal of Food Science, 54: 1302–1305.

McGuire, R, Agrawal, A.A. (2005). Trade-offs between the shade-avoidant response and plant resistance to herbivores? Tests with mutant *Cucumis sativus*. Functional Ecology, 19: 1025–1031.

Mody, K., Eichenberger, D, Dorn, S. (2009). Stress magnitude matters: different intensities of pulsed water stress produce non-monotic resistance responses of host plants to insect herbivores. Ecological Entomology, 34: 133–143.Munoz et al 2012

Muñoz, L.A., Cobos, A., Diaz, O. & Aguilera, J.M. Chia seeds: microstructure, mucilage extraction and hydration. Journal of Food Engineering, 108: 216–224.

Nobel, P.S., Geller, G.N., Kee, S.C, Zimmerman, A.D. (1986). Temperatures and thermal tolerances for cacti exposed to high temperatures near the soil surface. Plant, Cell, and Environment, 9: 279–287.

Nordenstam, B., Pelser, P.B. & Watson, L.E. (2009). The South African aquatic genus *Cadiscus* (Compositae-Senecioneae) sunk in *Senecio*. Comp. Newsletter, 47: 29–31.

Pan VS, McMunn M, Karban R, Goidell J, Weber MG, LoPresti EF. 2021. Mucilage binding to ground protects seeds of many plants from harvester ants: A functional investigation. Functional Ecology 35: 2448–2460.

Pan VS, Girvin C, LoPresti EF. 2022. Anchorage by seed mucilage prevents seed dislodgement in high surface flow: a mechanistic investigation. Annals of Botany 129: 817–830.

Parolin, P. (2005). Ombrohydrochory: rain operated seed dispersal in plants - With special regard to jet-action dispersal in Azioceae. Flora, 201: 511–518.

Pezzola, E., Mancuso, S. and Karban, R., 2017. Precipitation affects plant communication and defense. Ecology, 98:1693–1699.

Phan, J.L., Cowley, J.M., Neumann, K.A., Herliana, L., O’Donovan, L.A. and Burton, R.A., 2020. The novel features of Plantago ovata seed mucilage accumulation, storage and release. Scientific Reports, 10:1–14.

Phan, J.L. and Burton, R.A., 2018. New insights into the composition and structure of seed mucilage. Annual Plant Reviews Online, pp.63–104.

Rønsted, N., Chase, M.W. Abach, D.C., Bello, M.A. 2002. Phylogenetic relationships within Plantago (Plantaginaceae): Evidence from nuclear ribosomal ITS and plastid trnL-F sequence data. Botanical Journal of the Linnean Society 139: 323–338.

Ryding O. 2001. Myxocarpy in the Nepetoideae (Lamiaceae) with notes on myxodiaspory in general. Systematics and Geography of Plants 71: 503–514.

Tevis, L. 1958. Interrelations between the harvester ant Veromessor pergandei (Mayr) and some desert ephemerals. Ecology, 39: 695–704.

Teixeira, A., Iannetta, P., Binnie, K., Valentine, T.A. and Toorop, P., 2020. Myxospermous seed-mucilage quantity correlates with environmental gradients indicative of water-deficit stress: Plantago species as a model. Plant and Soil, 446: 343–356.

Vaughn, K.C., Bowling, A.J. and Ruel, K.J., 2011. The mechanism for explosive seed dispersal in Cardamine hirsuta (Brassicaceae). American Journal of Botany, 98: 1276–1285.

Villellas J, García MB. 2013. The role of the tolerance-fecundity trade-off in maintaining intraspecific seed trait variation in a widespread dimorphic herb. Plant Biology 15: 899–909.

Wagner, M.R, Mitchell-Olds, T. 2018. Plasticity of plant defense and its evolutionary implications in wild populations of Boechera stricta. Evolution, 72: 1034–1049.

Wardle DA, Ahmed M, Nicholson KS. 1991. Allelopathic influence of nodding thistle (Carduus nutans l) seeds on germination and radicle growth of pasture plants. New Zealand Journal of Agricultural Research 34: 185–191.

Western TL. 2012. The sticky tale of seed coat mucilages: production, genetics, and role in seed germination and dispersal. Seed Science Research 22: 1–25.

Whitford, W.G, Ettershank, G. 1975. Factors Affecting Foraging Activity in Chihuahuan Desert Harvester Ants. Environmental Entomology, 4: 689–696.

Yang X, Baskin CC, Baskin JM, et al. 2013. Hydrated mucilage reduces post-dispersal seed removal of a sand desert shrub by ants in a semiarid ecosystem. Oecologia 173: 1451–1458.

Yang, X., Baskin, C.C., Baskin, J.M., Guangzheng, L. & Huang, Z. (2012). Seed Mucilage Improves Seedling Emergence of a Sand Desert Shrub. PLoS ONE, 7: 1–9.

Yang X, Zhang W, Dong M, Boubriak I, Huang Z. 2011. The achene mucilage hydrated in desert dew assists seed cells in maintaining DNA integrity: Adaptive strategy of desert plant Artemisia sphaerocephala. PLoS ONE 6.

Zhou Z, Xing J, Zhao J, Liu L, Gu L, Lan H. 2021. The ecological roles of seed mucilage on germination of Lepidium perfoliatum, a desert herb with typical myxospermy in Xinjiang. Plant Growth Regulation in press. doi:10.1007/s10725-021-00702-y.

