## Supplement for "Drying conditions alter the defensive function of seed mucilage against granivores"

**Supplementary Table/Figure Legends:**

Supplementary Table 1: Species used in each study, as well as the source of the seed.

Supplementary Figure 1: Temperature dependence of 16 of the 22 species tested (all data is archived). The other species were either too weakly attached to measure reliably, we had too little seed to do a full temperature spread, or all treatments were not completed before SEW graduated.

Supplementary Figure 2: Survival of seeds exposed to harvester ants, when dried to dried to sandpaper at room temperature (Control), in a drying oven at 70C (Heat) and in vacuum desiccators with Drierite (Vacuum). Letters indicated treatments which differed significantly from each other in a post-hoc comparison of means.

Supplementary Figure 3: Attachment force of chia (*Salvia hispanica*) and cress (*Lepidium sativum*) on new and washed slides. The main effect of washing slide was not significant; however, the interactive effect of species and washing treatment was marginally significant (p=0.051). While we cannot say we conclusively found an effect, these results were suggestive enough of an effect that we decided to be conservative for the much larger data set and only used new slides for all force measurements.
